# Structural modelling and dynamics of the full-length Homer1 multimer

**DOI:** 10.1101/2025.05.26.655084

**Authors:** Zsófia E. Kálmán, András Czajlik, Brigitta Maruzs, Fanni Farkas, István Pap, Csilla Homonnay, Tomas Klumpler, Gyula Batta, Zoltán Gáspári, Bálint Péterfia

**Author notes:** equal contribution (Zsófia E. Kálmán, András Czajlik).

## Abstract

Homer proteins are modular scaffold molecules that constitute an integral part of the protein network within the postsynaptic density. Full-length Homer1 forms a large homotetramer via a long coiled coil region, and can interact with proline-rich target sequences with its globular EVH1 domain. Here we report an atomistic model of the full-length Homer1 tetramer along with the NMR solution structure of its EVH1 domain. Compared to the already available EVH1 structures, our NMR ensemble exhibits subtle differences, mostly in and around its partner binding region, suggesting the presence of ligand-induced conformational transitions. Molecular dynamics simulations of the long coiled coil reveal distinct regions with different stability and flexibility, with the N-terminal part of the coiled coil exhibiting the largest motions. Interestingly, this segment is highly conserved, pointing to the functional relevance of the observed dynamical features. Our results indicate previously unexplored aspects of the flexibility of the full-length Homer1 tetramer that might contribute to the dynamic rearrangements of the postsynaptic protein network linked to its functional transitions.

## Introduction

Excitatory synapses contain a highly organized network of different proteins. Beneath the postsynaptic membrane, a pronounced structure, called the postsynaptic density (PSD), is located [1], which is primarily organized by large multivalent scaffold proteins [2]. The three members of the Homer family (named Homer1, Homer2 and Homer3) are among the major scaffolding proteins in mammals [3] and Homer1 has several splice variants with different domain organization. The shortest isoform (Homer1a) contains only the N-terminal EVH1 domain and a short, disordered C-terminus, while the longer forms (Homer1b, Homer1c) both harbor the EVH1 domain connected to long C-terminal coiled coil segment by a long disordered hinge region [4, 5]. The coiled coil segment mediates higher-order assembly of multiple Homer polypeptide chains. M. K. Hayashi and colleagues proposed a model where the long coiled-coil region is divided into two segments: the first is a short, dimeric, parallel coiled coil, and a second starting as a parallel dimer continued as a tetramer by interconnecting the dimeric C-terminals in an antiparallel orientation [6] (Figure S1).

The highly conserved EVH1 domain belongs to the Class I type EVH1 domain that binds proline-rich short linear motifs (SLiMs) with the consensus sequences FPxoP and PPxxF (where x is any residue and o stands for hydrophobic residues) [7]. Most of the Homer interaction partners characterized so far bind to the EVH1 domain [8]. Multiple membrane receptors such as mGluR5 and IP3 receptors are linked to the deeper layer of the PSD through their interaction with Homer [9]. It also forms complexes with ion channels and transmembrane proteins such as TRPC1 [10]. Homer forms a mesh-like structure with Shank proteins and via this interaction it is also linked to the major scaffold protein, GKAP [6]. In addition, Homer1 is anchored to the cytoskeleton through the connection with drebrin and F-actin [4, 8]. It was also discovered that Homer contains an internal sequence SPLTP, called the P-motif, in the long hinge region between the EVH1 domain and the coiled coil segment. This motif has been suggested to form weak interactions with the EVH1 domain, although this has been observed only in the crystal state and not in solution, therefore the physiological relevance of this interaction is not yet clear [3].

Homer plays an essential role in determining the shape and density of the synapse, expected to be closely associated with the fundamental molecular mechanisms behind learning and memory. The presence of Homer1 isoforms seems to be tightly regulated as the short Homer1a isoform acts as a negative regulator blocking the formation of higher-order complexes of the larger, tetrameric isoforms. Homer1 is also a key protein in the reorganization of the PSD during sleep, when the dimeric Homer1a isoform gets dominant over the larger tetrameric assembly and thus disrupts the direct connections between mGluR5 receptor subunits in the postsynaptic membrane and the IP3 receptors in the ER [9]. The dysfunction of Homer proteins was also linked to many neurological symptoms, Homer is believed to play a role in schizophrenia, Fragile X syndrome and many more [8].

In this work we describe a structural model of the full-length Homer1c isoform by combining the globular, intrinsically disordered and coiled-coil segments into multimeric structures. A key aspect is to describe the local and global flexibility of the protein. To this end, we have determined the solution structure of the EVH1 domain by NMR spectroscopy and have performed molecular dynamics simulations on the dimeric and tetrameric complex of Homer.

## Materials and Methods

### Cloning, protein expression and purification

*Mus musculus* Homer1 EVH1 domain coding insert was ligated into the Nde1 and BamH1 restriction sites of a modified pET-15b plasmid backbone (Novagen) - that contains an N-terminal 6xHis-tag and a tobacco etch virus (TEV) protease cleavage site. The expressed protein corresponds to UniProt Q9Z2Y3, 1-118 sequence: GSHMGEQPIFSTRAHVFQIDPNTKKNWVPTSKHAVTVSYFYDSTRNVYRIISLDGSKAIINSTITPNMTFTKTSQKFGQWADSRANTVYGLGFSSEHHLSKFAEKFQEFKEAARLAKEKSQ where the first 3 amino acids remain after cleavage of the expression tag. Note that Mus musculus and Human sequences are identical at this section of Homer1 protein. The recombinant protein was expressed in BL21 (DE3) *Escherichia coli* bacterial cells (Novagen) for 20 hours incubation at 20 °C with 150 rpm shaking after induction with 1mM IPTG (isopropyl β-D-1-thiogalactopyranoside) (Sigma) at a cell density of 4 MFU. After centrifugation, the cell pellets were stored at -20 °C until further use. Unlabeled EVH1 protein was expressed in cells grown in LB medium. For isotopically labeled (^15^N and ^15^N, ^13^C) protein production M9 medium was used [11] supplemented with 0.2x Trace element mix [12], 2.5 g/L ^15^NH_4_Cl (Cambridge Isotope Laboratories, Cambridge, MA), and 4 g/L unlabeled glucose or [^13^C]-D-glucose in the case of double labeled version. Harvested cells were disrupted by sonication in a 10% suspension of lysis buffer (300 mM NaCl, 50 mM sodium phosphate, pH 7.4) using an ultrasonic homogenizator (BioLogics). Homer EVH1 proteins were purified from the supernatant after homogenization with a 5 ml Bio-Scale™ Nuvia™ IMAC Ni-affinity column (Bio-Rad) equilibrated with binding buffer (50 mM sodium phosphate, 300 mM NaCl, pH 7.4). This binding buffer was also used for washing and elution steps, supplemented with 50 mM and 250 mM of imidazole, respectively. IMAC purification was followed by His-tag cleavage with TEV protease (25 ug TEV/g EVH1, 10 °C, ON). After TEV treatment, the buffer was exchanged to low salt phosphate buffer (binding buffer with 20 mM NaCl) using HiTrap desalting columns with Sephadex G-25 resin (Cytiva), and the sample was further purified by ion exchange chromatography, using a Bio-Scale™ Mini Macro-Prep® High S column. Gradient elution was performed by adding NaCl to the binding buffer up to 500 mM. Reverse IMAC was applied to get rid of the cleaved His-tag and the TEV enzyme. In case of samples for NMR measurement an additional SEC (size exclusion chromatography) step was included using a SEC70 analytical gel chromatography column (Bio-Rad). Before and after SEC, the sample was concentrated with the Amicon® Ultra 15 ml Centrifugal Filter unit with 3 kDa MWCO. The recombinant EVH1 protein was eluted in low salt sodium phosphate buffer (50 mM sodium phosphate; 20 mM NaCl, 0.02% NaN_3_; pH 7.4). To determine its purity and molecular weight, SDS-PAGE (sodium dodecyl sulfate - polyacrylamide gel electrophoresis) was applied, and the concentration was measured by determining A280 using a NanoDrop2000 instrument and the extinction coefficient estimated from the protein sequence by a protein calculator software (Sourcefolge:https://protcalc.sourceforge.net/).

### Far-UV CD spectroscopy

Far-UV CD spectroscopy measurement was performed using a JASCO J-1500 CD spectrometer (JASCO Corporation, Tokyo, Japan) in 195-250 nm spectral range, 50 nm/min scanning speed, 1 nm bandwidth, 0.2 nm step size, 0.5 s response time and 3 scans of accumulation and baseline correction. It was recorded in a 1 mm pathlength quartz cuvette (J/21, Jasco). Sample concentrations were 10 μM in 50 mM Sodium Phosphate Buffer, 20 mM NaCl, pH 7.4 buffer. In order to check temperature dependence, the 20 °C-60 °C interval was used, measurement was done in every tenth degree.

Secondary structure calculation and data evaluation was performed with BeStSel (https://bestsel.elte.hu [13])

### Mass Spectrometry

To determine the exact molecular weight, mass spectrometry measurements were performed on the TEV-treated H118 protein. HPLC-MS measurements of the proteins were performed using a Shimadzu LC-MS-2020 device with a Reprospher-100 C18 column and a positive-negative double ion source (DUIS±) with a quadrupole MS analyzer in a range of m/z 50-1000. Gradient elution was applied using eluent A (0.1% formic acid in water) and eluent B (0.1% formic acid in acetonitrile). According to the MS results, the efficiency of isotopic labeling was found to be 98% for ^15^N and 97% for ^13^C labelling (Table S1).

### NMR Measurement conditions and data analysis

For data collection, samples of 600 µl containing 190 µM ^13^C, ^15^N-labeled protein in a solution of 50 mM Sodium Phosphate Buffer, 20 mM NaCl, 0.02% NaN_3_ at pH 7.36 were prepared. Measurements were performed on a Bruker Avance NEO 700 MHz spectrometer at 298 K. For resonance assignment, standard 2D (^1^H-^15^N HSQC, ^1^H-^13^C HSQC) and 3D triple-resonance (HNCA, HN(CO)CA HNCACB, HN(CO)CACB, HNCO, HN(CA)CO, HCCONH, HCCH-TOCSY) and HBCBCGCDHD spectra were acquired. To aid structure determination, 2D NOESY, 3D TOCSY-HSQC, and NOESY-HSQC spectra were also recorded.

Initial structure calculation was performed with the ATNOS/CANDID/Cyana suite using 7 cycles. For the calculation, NOE restraints along with TALOSN-based [14] dihedral angle restraints were used. To improve local geometry, the 20 best structures obtained were further refined with XPLOR-NIH using the NOE restraints only. For this, a gentle simulated annealing protocol was applied with an upper temperature of 150 °C to preserve the input conformations while allowing optimization of the local geometry. Before refinement, NOE restraints were manually curated to exclude any remaining non-trivial redundancies like both specific and ambiguous inclusion of geminal protons in separate restraints to the same partner atoms. The stereospecific restraints deduced by the initial protocol were retained, and atom names were converted between the Cyana [15] and XPLOR [16] formats with in-house scripts. As the initial ATNOS/CANDIS/Cyana protocol uses the so-called sum averaging method, the restraints were rescaled and binned to be suitable for the simple distance averaging setting used in our XPLOR-NIH refinement step.

Visualization of the structure was performed using MOLMOL [17] and Chimera [18]. Electrostatic surface calculation was performed using ABPS as available through the site https://server.poissonboltzmann.org/ [19].

### Small Angle X-ray scattering measurements

For SAXS measurement the sample preparation protocol was similar to the above mentioned NMR sample preparation. The only difference was that unlabeled proteins were used. SAXS data was collected from 8, 4, 2 and 1 mg/ml protein samples.

SAXS data was collected using a Rigaku BioSAXS-2000 instrument at CEITEC (Brno, Czech Republic) equipped with a HyPix-3000 detector at a sample-detector distance of 0.48 m. Scattered intensity was measured in range 0.008-0.65 1/Å, where q= 4π sin θ/λ; 2θ is the scattering angle and λ = 0.154 nm. Six 10 minute frames were collected at a sample temperature of 25 °C. The data were normalized to the intensity of the transmitted beam and radially averaged. Scattering curves from individual frames were checked for radiation damage and averaged. The corresponding scattering from the solvent-blank was subtracted to produce the scattering profile.

Structure-based calculation of SAXS curves was performed with Pepsi-SAXS [20]. The per-conformer curves were averaged.

### Model building and MD simulations

For the Homer tetramer modeling, the solved rat crystal structures covering the protein terminals (1ddv [21] for the N-terminal EVH1 domain and 3cve [6] for the C-terminal tetramer CC) were complemented by building the intermediate regions. The disordered-CC boundary was defined using MobiDB [22]. Disordered segments were modelled using Dipend [23] and Modeller in Chimera [18]. The general parameters of the coiled coil segment was analysed using CCBuilder [24] and the refinement of the parameters and final modelling using Isambard [25]. The structure was tested with SOCKET [26] and minimized using FoldX [27].

Computations were performed on the Komondor supercomputer. The models were simulated with GROMACS [28] using explicit solvent classical MD. AMBER99SB-ILDN protein force field was used; the box setting was a minimum distance of 1.0 n. For the initial model five parallel 200 ns runs for the dimers were carried out (md_200_1, md_200_2, md_200_3, md_200_4, md_200_5).

For further analysis we used different tools. Secondary structure was predicted using DSSP [29]. From the DSSP predictions we also calculated the relative surface accessible area. SOCKET [26] was used to assign coiled coil regions and heptad positions to the structures. Structures were visualised using Chimera [18]. We both used ColabFold [30] and AlphaFold3 [31] for the additional modelling. Ca-distances, salt bridges were calculated with in-house Perl and Python scripts. Matplotlib was used for visualization. Sequence alignment was carried out using Jalview [32]. Vertebrate orthologous sequences were downloaded from OMA [30] and Shannonentropy was calculated with an in-house script.

## Results

### Expression of the murine Homer1 EVH1 construct

To characterize the EVH1 domain, the segment 1-118 from *Mus musculus* Homer1 (UniProt ID: Q9Z2Y3) was selected. Murine, rat and human sequences are 100% identical at this region of the Homer1 protein (Figure S2). The molecular weight (MW) of the unlabelled EVH1 domain was calculated to be 13.7 kDa (13736.38 Da) based on the encoded amino acid sequence. Mass spectrometry (MS) measurements confirmed that the correct sample was produced (13718.37 Da) and SDS−polyacrylamide gel electrophoresis (SDS−PAGE) also supports this result (Figure S3). Far-UV CD spectroscopy analysis of the successfully expressed Homer1 EVH1 domain in 50 mM Sodium Phosphate Buffer, 20 mM NaCl at pH 7.4 was consistent with an intact globular structure dominated by β-sheets (Figure1A). BestSel [13] analysis indicates 13.6% alpha-helix, 46.3% beta-sheet and 40.1% non-classified secondary structure elements. Temperature-dependent CD measurements do not indicate major structural changes between 25 to 60 °C. Thus, the expressed EVH1 domain has a well defined, stable structure (Figure 1A).

**Figure 1.**
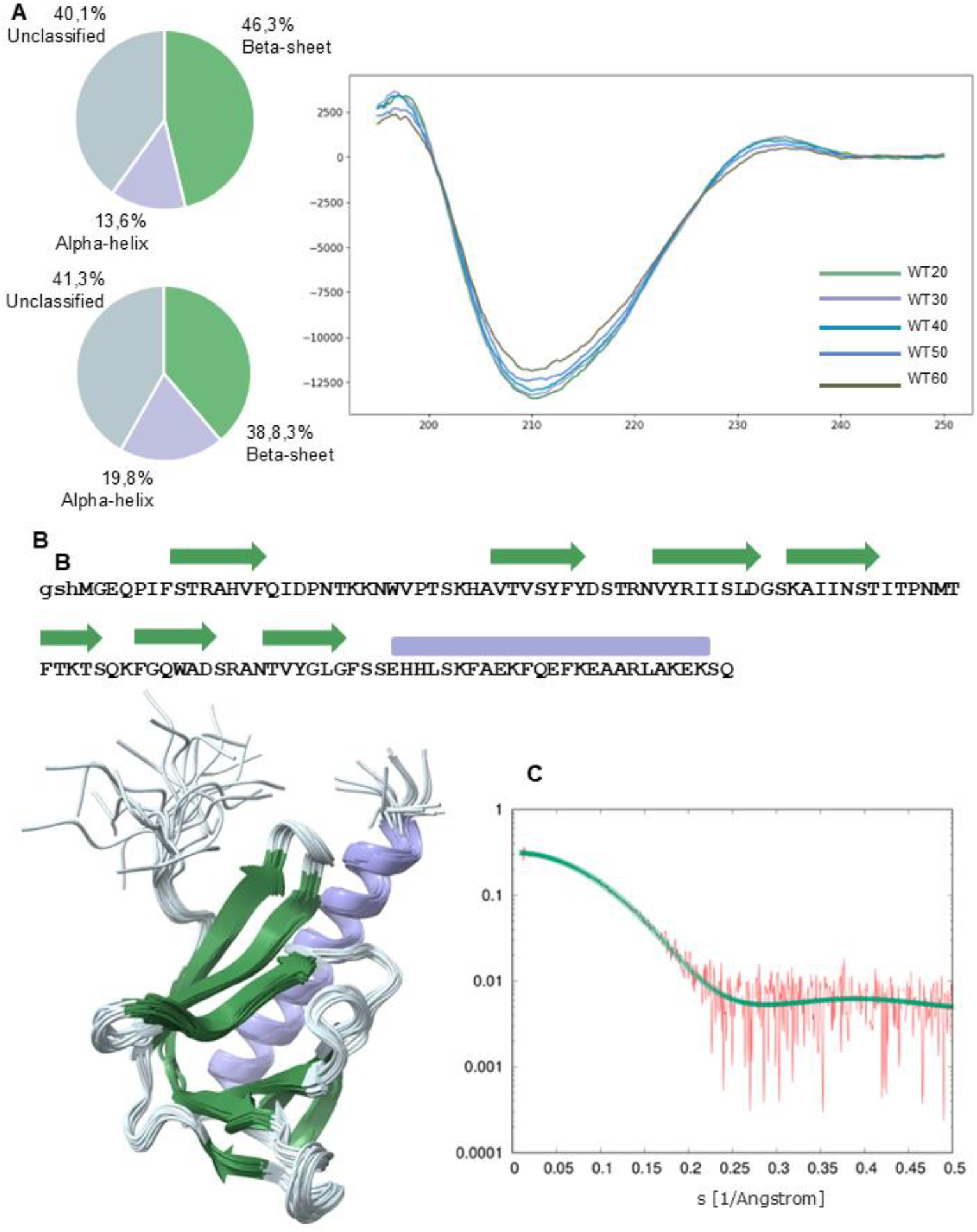
General features of the EVH1 domain (A) Beta-sheet dominated globular structure of the EVH1 domain. CD spectra of the EVH1 domain construct. Secondary structure percentages calculated from the CD spectra using BestSel (upper diagram) from the average structure using DSSP (bottom diagram). (B) Position of the secondary structural elements in the sequence and in the structure. Green: Beta sheet, Purple: Alpha-helix, Grey: Linkers and loops. (C) Comparison of the measured (red) SAXS curve and the one calculated and averaged for the 20 conformers.

### Solution structure of the EVH1 domain

The ^1^H-^15^N HSQC spectrum of the EVH1 construct reveals a well-folded structure, characterized by high signal dispersion and predominantly well-resolved peaks. Complete backbone and partial sidechain NMR chemical shift assignments of the EVH1^1-118^ domain were obtained. For the backbone, 99% of N, 95% of H and 99% of C atoms, whereas for the side chains, 86% of C and 88% of H atoms could be assigned. Chemical shifts have been deposited in BMRB under ID 34990 and the coordinates in the Protein Data Bank under ID 9QUX (Figure S4).

The solution structure of the domain exhibits high overall similarity to the other available EVH1 structures, forming part of the larger PH domain family [7]. Throughout the text, we will use residue numbering matching the UniProt sequence, relative to which the numbering in the PDB entry is shifted by 3 residues because of the presence of the N-terminal expression tag. The fold can be described as a β-sandwich formed by 7 strands, flanked by a long C-terminal α-helix. The first β-strand (strand A) is largely bent, its N-terminal region pairs with strand B, whereas its C-terminal part with strand G, therefore, the fold is also sometimes described as a β-barrel [34]. In the solution structure of Homer1 EVH1, the seven β-strands are formed by residues 8-15, 32-39, 44-51, 54-60, 67-71, 74-79, 84-89, as identified by DSSPcont [35] (Figure1B, Figure S5). Most of the consecutive strands are linked by short turns, and even the longer, more irregular intra-strand segments 16-31 and 61-66 contain turn regions identified by DSSPcont. Residues 90-92, linking the last β-strand to the α-helix form a bend. The α-helix extends from residue 93 to 115, thus, only the last three residues at the C-terminus of the construct do not adopt helical conformation (Table S2). All buried amide NH groups participate in hydrogen bonds. SAXS measurements are in good agreement with the calculated structure, with a X^2^ of 1.067 between the experimental and the averaged back-calculated curve for the 20 conformers (Figure 1C). The SAXS data were deposited at SASBDB under the accession code: SASDXK2.

The upfield chemical shifts of the sidechain atoms of Lys 102 are indicative of the proximity of an aromatic ring, consistent with the calculated structure where this residue is surrounded by the side chains of Phe7, Tyr36, Phe99 and Phe106 (Figure 2A). Phe99, Lys102 and Phe106 are located in the long α-helix, whereas Phe7 and Phe36 are in β-strands 1 and 2, respectively. Thus, this cation-pi interaction links the C-terminal helix to the first half of the β-sheet. It can be noted that in our solution structure, the orientation of Lys102 is different from that observable in the Homer1 EVH1 X-ray structures, its side chain being closer to the aromatic rings in the NMR ensemble.

**Figure 2.**
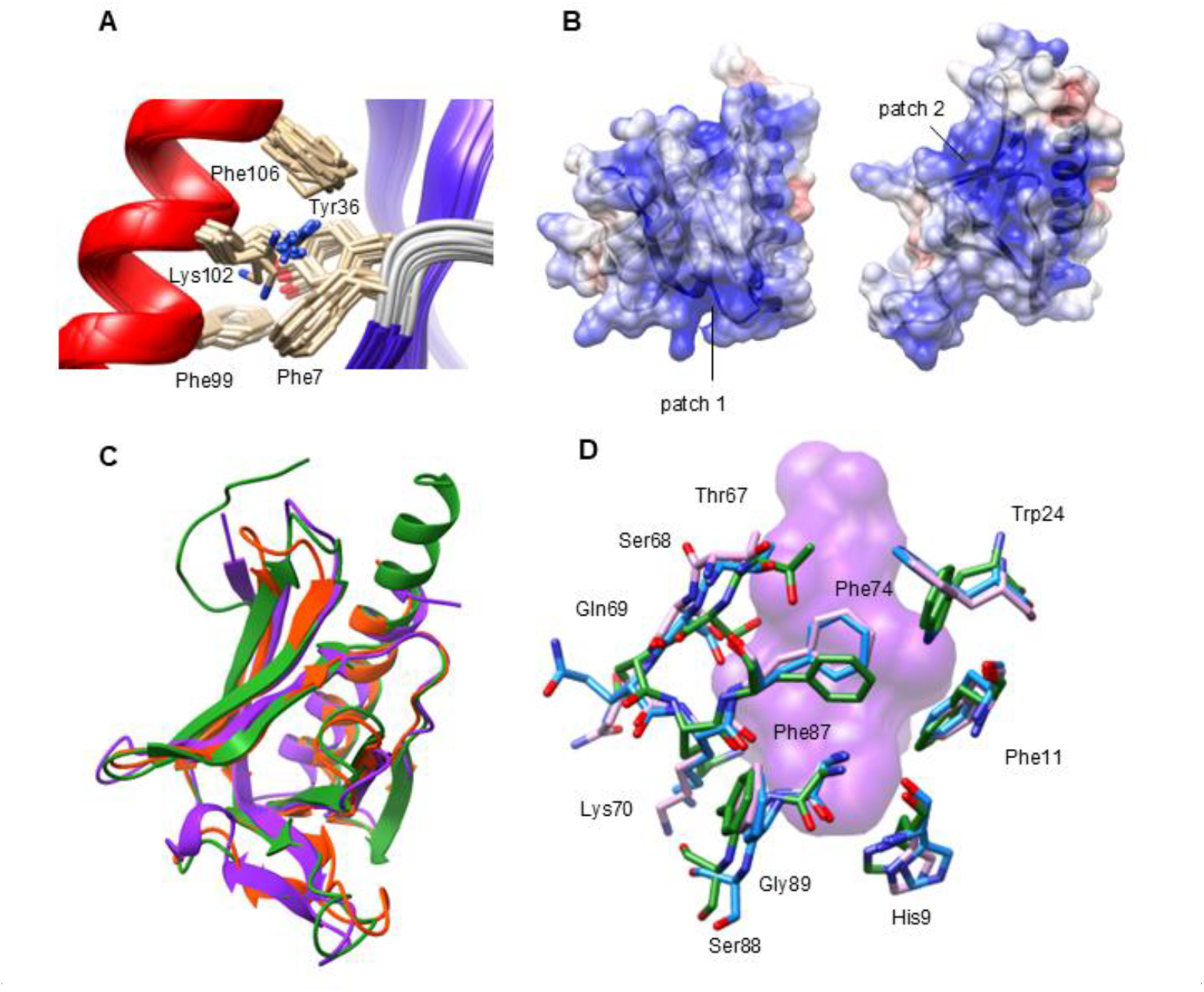
Local features of the EVH1 structure. A) The putative cation-pi interaction between Lys102, Phe7, Tyr36, Phe99 and Phe106 in the NMR ensemble. B) Electrostatic surface of the Homer1 EVH1, illustrated using the medoid model in the ensemble. Blue shades depict positive, red shades negative surface areas. C) comparison of EVH1 domains. Green: the NMR ensemble described in the present study. Orange red: other available Homer1 EVH1 structure. Purple: additional EVH1 structure. D) Orientation of the ligand-binding residues in the medoid structure of our NMR ensemble (color by atom type), the structure 1ddw (without ligand, blue) and 1ddv (liganded, purple). The position of the ligand in the entry 1ddv (MGluR peptide) is shown with a semi-transparent purple surface.

The electrostatic surface of the EVH1 domain is predominantly positive, with two regions with high positive charge. The first one (patch 1) is located between the first β-strand and the N-terminus of the helix, close to the ligand-binding site, and the second (patch 2) between the C-terminal segment of the helix and the tip of the β-barrel (Figure 2B).

Compared with other available Homer EVH1 structures (1ddv, 1ddw, 1i2h, 1i7a, 2p8v, 5zz9 [21, 3, 36, 37, 38]), the largest difference can be observed in the loop regions between residues 17-29 and 40-43. The former, longer loop forms the tip of a beta-hairpin that contributes a number of residues to the partner binding site (Figure 2C). In the majority of Homer1 EVH1 crystal structures, the conformation of this loop is highly similar, regardless of whether they are in the free or in a bound state (Figure 2C, Figure S6, Table S3). According to analysis with ePISA [39], this loop participates in crystal contacts with neighboring molecules, providing a possible explanation of this observation. The only Homer1 X-ray structure with a different loop conformation is 1i2h, which represents an EVH1 domain binding to the P-motif (SPLTP) of neighboring Homer1 molecules in the crystal [3]. Considering other available EVH1 domain structures, this region shows yet again different conformations in Enabled, N-Wasp and Spred2 proteins, thus, we speculate that this loop might undergo minor rearrangements upon partner binding. In addition, in our NMR ensemble the indole ring of Trp24 displays an orientation that is almost perpendicular to those observed in the other EVH1 structures (Figure 2D). Trp24 is one of the key residues involved in partner binding, and the reorientation of its side chain might well accompany the binding of the partner Pro-rich sequences as the aromatic cluster of Phe14, Trp24 and Phe74 reorganizes to accommodate the pyrrolidine rings.

### Modeling the coiled coil region

To model the full-length Homer1 (Homer1c isoform), we used the previously available structures of the globular EVH1 domain (PDB ID: 1ddv [21]) and the C-terminal fragment forming a four-stranded antiparallel coiled coil (PDB ID: 3cve [6]) (Figure S7). Since both the N- and C-terminal regions are only determined for the *Rattus norvegicus* protein, we chose to model the full-length Homer1 of this species. The sequential differences between rat, mouse and human is shown in Figure S2. The characteristic parameters (radius, pitch, interface angle) of the tetramer structure were determined. Based on these characteristics, the long coiled-coil regions were modelled first with CCbuilder [24] and later with Isambard [25] exploring different settings. The parallel long chains were aligned to the tetramer and inspected visually. Only residues 112-189, linking the EVH1 domain to the coiled coil region, were modeled explicitly as disordered (see Methods). It is expected that there are other fully or partially disordered regions within the dimeric coiled coil region or at the transitional structure between the two- and four-stranded coiled coil segments. However, we decided against including any such intermediate disordered regions in the initial model as their boundaries could not be precisely predicted. The challenges of modeling a long coiled coil with alternating regular and unfolded segments include proper linking of the different regions while maintaining regularity in the superhelical segments, which could introduce unrealistic structural features that could be hard to correct at later stages. Our working hypothesis was that flexible regions modeled as coiled coils will turn out to be unstable during the molecular dynamics simulations, providing a better estimate of the actual dynamics of the region than those deducible from sequence-based predictions only.

### Molecular dynamics simulation of the dimeric and tetrameric form of full-length Homer1

Multiple rounds of all-atom molecular dynamics simulations in explicit water were carried out for both the dimer form and the tetramer (more than 11 000 atoms) for 200 ns per run. For the dimeric structure, five parallel runs with different random initial velocities were used and will be referred to as dimer_1-dimer_5. The simulated trajectories were converted to 20 000 models/run for the dimer and 1000 models for the tetramer. The parallels capture quite different dynamics and conformations as seen from the calculated RMSDs (Figure S8).

Analysis of the secondary structure and the formation of knobs-into-holes interactions, characteristic of coiled coils, were performed using DSSP [29] and SOCKET [26], respectively. This analysis showed that the stability of the superhelical structure, as expected, is not uniform along the modeled coiled coil (Figure 3). It is important to note that there are several substantial differences observed between the simulation trajectories, therefore, we will focus on the commonly occurring features unless noted otherwise. Considering the consensus of the five MD runs, two stable regions can be identified, one covering residues 260-270 and the other spanning residues 300-325. In one of the simulations (dimer_1), the whole C-terminal segment of the coiled-coil from residue 250 remains stable (Table S4).This observation seems to be largely independent from the initial conformation as we run a shorter simulation on an AlphaFold predicted dimer (Figure S9) and these regions showed similar tendencies regarding flexibility. The distribution of intermolecular salt bridges fit well with the stability data if we consider heptad positions. There are five possible pairs formed between residues in heptad position ‘e’ in one chain and ‘g’ in the other. These pairs are located in positions 258-260, 302-307, 309-314, 321-323 and they can be consistently identified in each simulation (Figure 3). Outside of this region (residues 190-250) we did not observe any straightforward relationships between coiled coil stability, assigned heptad positions and possible salt bridges, indicating a more complex dynamic behavior of the N-terminal part of the long coiled coil. Interestingly, intramolecular salt bridge formation shows substantial variability in the different runs especially in the N-terminal region of the coiled-coil (residue 190-270). The region 270-295 exhibits different interaction patterns in the individual runs, but only maintains the coiled coil conformation during the dimer_1 simulation.

**Figure 3.**
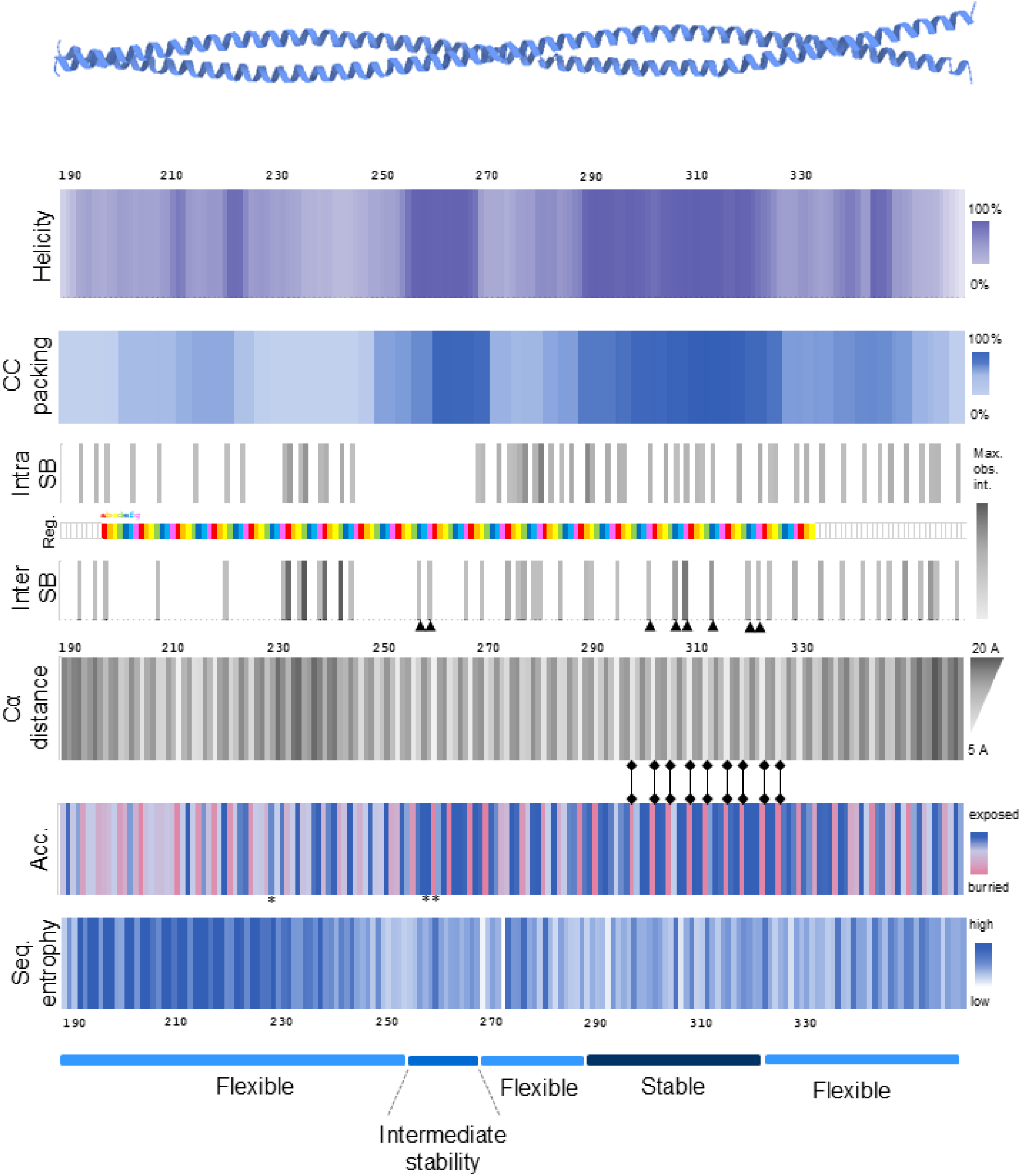
Analysis of coiled-coil stability. Different panels show the aspects of the observed differences in the structural stability along the coiled-coil. The average of the five parallel runs was calculated and plotted. Panels: i) Predicted stable helical segments during the simulations are in shade of dark purple (DSSP [26]). ii) Predicted stable coiled coil segments during the simulations are in shade of dark blue (SOCKET [23]) iii) Possible intrachain salt bridge formation, the darker gray shows more detected interactions iv) Heptad positions along the coiled-coil(Coloring according SOCKET-defined heptad positions: ‘a’-red, ‘b’-orange, ‘c’-yellow, ‘d’-green, ‘e’-dark blue, ‘f’-light blue,’g’-pink) v) Possible interchain salt bridge formation; arrows indicating, e’ and, g’ positions in the heptade repeat vi) Average Ca alpha distances between the two coiled-coil chains vii) Average accessibility of the position during simulation. For the C-terminal segment several amino acids in position, a’ and, d’ show regular patterns in vi) and vii). viii) Shannon entropy in vertebrates the more conserved positions are the darker. ix.) Five distinguishable regions of the coiled-coil segment (Numerical data is available in Table S4).

Analysis of Cα-Cα distances between identical residues in the interacting chains forming the coiled coil offers another means to assess the flexibility of the structure. These distances exhibit a periodicity according to the positions in the heptad repeats, but also show variability for residues in the same positions but in different regions, indicating non-uniform stability along the superhelical structure. The observed enlarged Cα-Cα distances are consistent with the results of the DSSP [29] and SOCKET [26] analysis and also indicate partial local unfolding of the coiled coil (Figure 3). There is an extremely regular segment of the coiled coil aligning with the second stable region (residue 300-325) where Cα-Cα distances are extremely similar for each run and consistent with the buried nature of residues in positions ‘a’ and ‘d’. This segment also shows a highly regular pattern of buried and exposed residues - in accordance with their position in the heptad (Figure 3). We investigated whether the structurally more stable segments show higher evolutionary conservation at the sequence level. Shannon-entropy calculated for the individual residue positions shows surprisingly high conservation at the N-terminal region of the coiled coil that proved to be the most unstable during our simulations (Figure 3).

The dimeric Homer1 models exhibit high structural flexibility resulting in different kinds of structural rearrangements leading to different global shape of the assembly and different relative position and orientation of the EVH1 domains (Figure 4). We have analyzed three aspects of such structural features: a) the distance between the two EVH1 domains b) presence of kinks within the coiled-coil c) the rotation of the two binding pockets in the EVH1 domains relative to each other. The distance between the two EVH1 domains, measured between the positions calculated from the average of the domain can vary substantially from over maximum 200 Å to a minimum distance of 40 Å (Figure S10A). This distance decreases with time during our molecular dynamics simulations. This certainly does not adequately describe the cellular behaviour of the system, but it highlights the large distances that even such a dimeric complex can bridge, and also shows (in the case of dimer_3 and dimer_5, most notably in the 0-100 ns time scale) how dynamically this distance can evolve (Figure S10B). The rotation of EVH1 domains with respect to each other is highly heterogeneous for different models. The long disordered region provides a very high degree of flexibility in the spatial location of the domains, which can be important for binding to different partners (Figure S10C). The motion resulting from the kinking of the coiled coil reflects its structural flexibility in the dimeric state in comparison to the tetramer that seems to be more rigid in this manner (Figure S10D).

**Figure 4.**
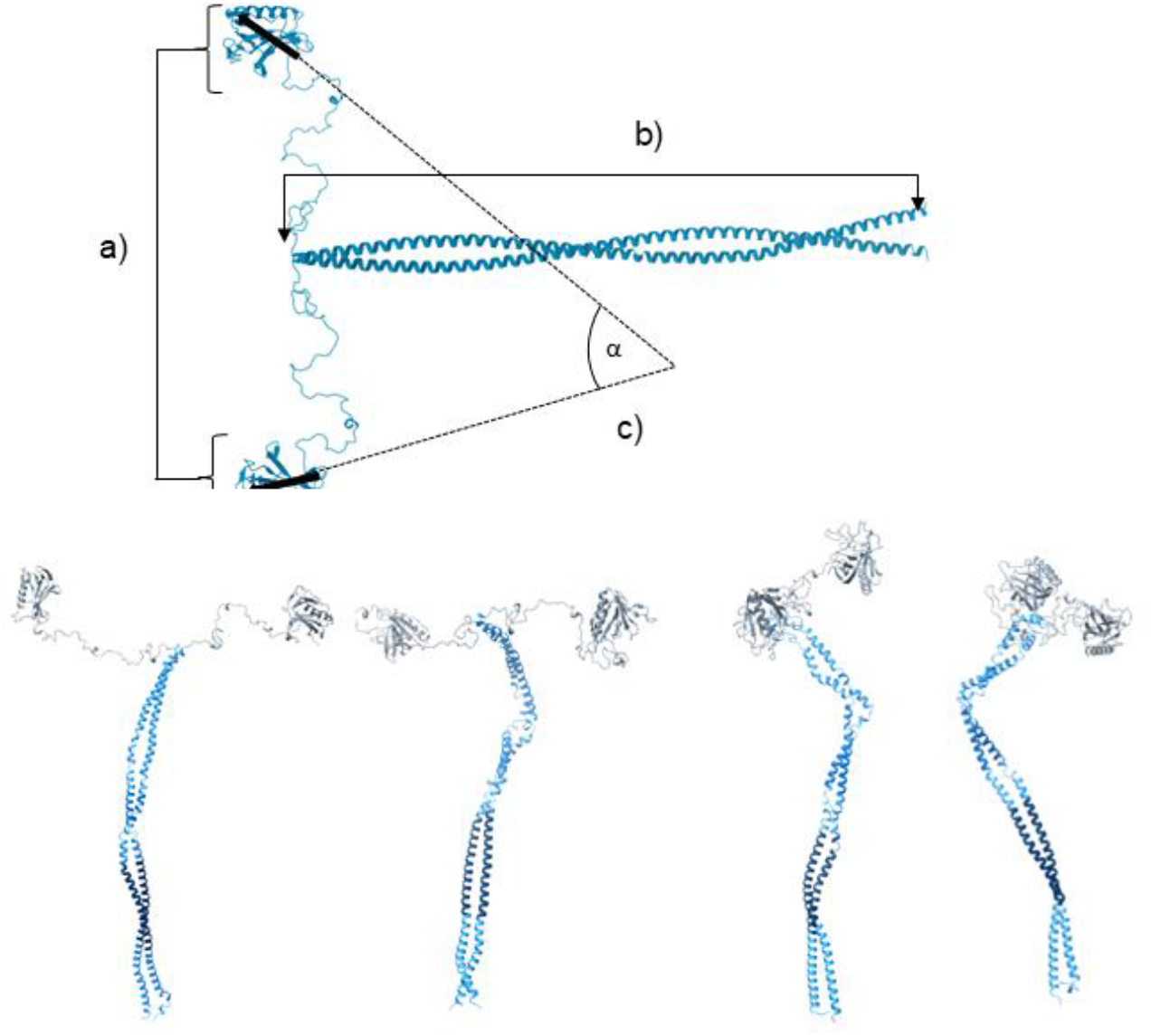
The analysed features of the flexibility shown of the structure a) distance of the average position in the domain b) distance of the N and C-terminal of the coiled-coil c) rotation of the two binding pockets in the EVH1 domains relative to each other in 3D. Structures representing the structural flexibility of the dimeric Homer1. Coloring according to five the regions with different stability as depicted in Figure3 panel ix.

Based on an interaction observed in the crystalline state, Homer1 EVH1 was previously suggested to bind to its internal P-motif, although this could not be confirmed in solution [3]. Nevertheless, the presence of such an interaction can not be readily dismissed in a crowded environment and, especially, in the context of a multimeric protein. Therefore we have investigated whether this interaction could occur in the context of the tetrameric form either within a single Homer1 chain or between neighboring chains in the complex. From our initial modelling it seems that the formation of an intrachain interaction between the P-motif and the EVH1 domain is sterically possible, however the small helical fragment at the C-terminal of the EVH1 domain should not be longer than 15 residues (Figure S11). There are also variations in the length of the helix in the available crystal structures.

## Discussion

In this study we have explored the structure of the full-length Homer1 tetramer, using NMR spectroscopy to provide the first solution structure of the globular EVH1 domain and applying modeling and molecular dynamics for the characterization of the coiled coil region. The solution structure of the EVH1 domain aligns well with the overall fold of previously determined EVH1 structures, but also suggests the presence of subtle structural rearrangements upon ligand binding. The local conformation around the ligand-binding region, especially residues 18-21, forming a turn after the first beta strand, is different from that observed in the available Homer EVH1 X-ray structures. On the other hand, this region exhibits variability when considering a larger set of EVH1 domains, lending support to the hypothesis that this segment can undergo functionally relevant motions The possible functional relevance of such rearrangements, as well as of the positively charged surface patches, especially that of patch 2, located relatively far from the described ligand binding site, remain to be explored.

To explore the architecture of the full Homer1 tetramer, we have applied a complex pipeline using tools specifically designed for coiled coil modeling and analysis. We have used the available X-ray structure of the tetramerization region and fitted a model of the long dimeric coiled coil to it. The resulting models, containing either a dimeric segment or the full tetrameric assembly, were analyzed to assess its dynamics and stability features. Our molecular dynamics simulations indicate the presence of segments with different stability and flexibility in the long coiled coil region, with the most stable region being the C-terminal segment next to the tetramerization domain. The increased flexibility towards the N-terminal part of the coiled coil makes it possible for the full tetramer to adopt various global shapes besides the fully extended linear one. This kind of flexibility in long coiled coil segments has been observed for other proteins both with experimental and theoretical methods. The intramolecular coiled-coil of Rad50 also showed irregular flexibility in their long coiled-coil segment [40]. Tropomyosin coiled-coil flexibility is also required for its function [41]. The kink formation was revealed for laminin supported by experimental data measured with atomic force microscopy and also observed in coarse-grained MD simulations [42].

The residue conservation pattern of Homer1 segments does not correspond to the location of stable and unstable coiled coil segments identified by our simulations. However, it must be noted that conservation indicates functional importance and not necessarily structural stability. Thus, it can be speculated that the observed conservation features indicate that a precisely tuned flexibility is necessary to maintain Homer1 function. Moreover, the higher flexibility of this region relative to the C-terminal part does not exclude its role in initiating multimerization, as is currently proposed for cotranslational assembly of protein complexes [43], and is well in line with our earlier observations on the importance of such regions based on the pattern of germline mutations [44].

The presence of kinked structures within the coiled coil and the flexibility of the disordered linker enable a wide range of relative distances and orientations for the four EVH1 domains in the tetramer. Thus, the full-length Homer might not only bridge large distances, but could also provide a permanent physical link between partner molecules and complexes during larger structural rearrangements within the postsynaptic protein network. The role Homer fulfills in synapse might be essential, the EVH1 domain is highly conserved, even the Drosophila sequence has remarkable identity to the human protein. The coiled coil segment has also changed relatively little in vertebrates. It might also be speculated that the EVH1 domain might not be the only region that can participate in various protein:protein interactions, and these additional interactions might play a role in modulating the flexibility of the coiled coil rod. The best candidate for such an interaction is the P-motif, capable of forming complexes with the EVH1 domain in the crystalline state, although not in solution [3]. Still, its conservation within a disordered segment is best explained by its role as a potential binding motif, and it might also be possible that the same applies to the conserved yet flexible N-terminal part of the long coiled coil. Our results altogether suggest that Homer1 does not only act as a passive linker between its partners but might have more complex roles in the organization of the postsynaptic protein network.

## Acknowledgements

This work was supported by the Hungarian National Research, Development and Innovation Office (NKFIH) (grants, OTKA K 137947, TKP2021-EGA-42, and 2021-4.1.2-NEMZ_KI-2022-00027). We acknowledge the Digital Government Development and Project Management Ltd. for awarding us access to the Komondor HPC facility based in Hungary.

The authors are grateful to Gitta Schlosser for the HPLC-MS measurements, and to Viktor Farkas for the supervision of CD spectroscopy measurements. We acknowledge CF Biomolecular Interactions and Crystallography of CIISB, Instruct-CZ Centre, supported by MEYS CR (LM2023042) and European Regional Development Fund-Project,, Innovation of Czech Infrastructure for Integrative Structural Biology” (No. CZ.02.01.01/00/23_015/0008175)

## Data availability

Nuclear magnetic resonance data for the structural ensemble reported in this article have been deposited at the Biological Magnetic Resonance Data Bank, under deposition ID:34990. The NMR structure of the EVH1 domain is deposited in PDB and accessible under the ID:9QUX.The corresponding small-angle X-ray scattering data have been deposited to the Small Angle Scattering Biological Data Bank with the ID: SASDXK2. The modelled Homer1 teramer is available in ModellArchive with the following ID: ma-cp0vl.

## Author contributions

**Zsófia E. Kálmán**: investigation, formal analysis, writing – original draft, visualization; **András Czajlik**: investigation, formal analysis; **Brigitta Maruzs**: investigation, formal analysis; **Fanni Farkas**: investigation, formal analysis, writing – original draft; **István Pap**: investigation; **Csilla Homonnay**: investigation; **Tomas Klumpler**: investigation, writing – original draft; **Gyula Batta**: investigation, conceptualization; **Zoltán Gáspári**: investigation, conceptualization, writing – original draft, visualization, supervision, funding acquisition; **Bálint Péterfia**: investigation, conceptualization, writing – review and editing, supervision

