## Supplemental Figures 1-11 for "Structural modelling and dynamics of the full-length Homer1 multimer"

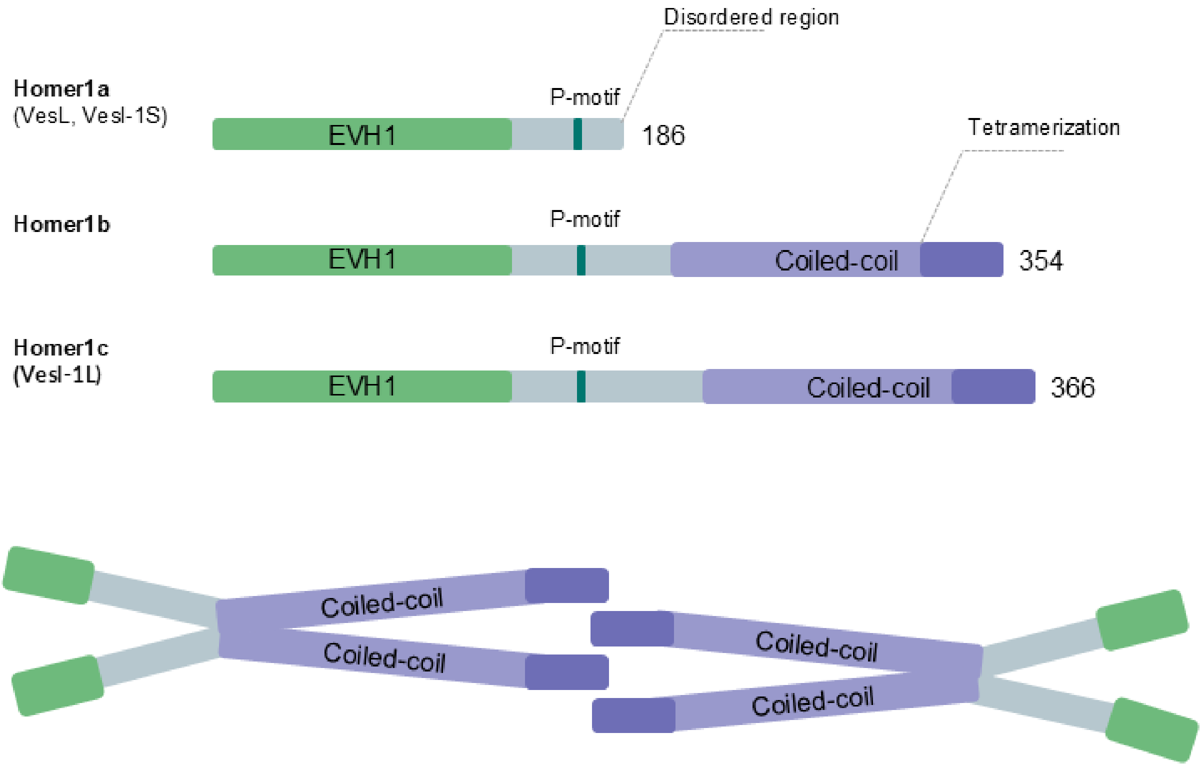


**Figure S1** Top: Isoforms of murine Homer1 proteins. Bottom: Schematic figure on the tetramerization of Homer1c. Experimental structures are only available for the EVH1 domain (see Table S3) and the tetrameric coiled coil region (PDB: 3CVE) [4,5].

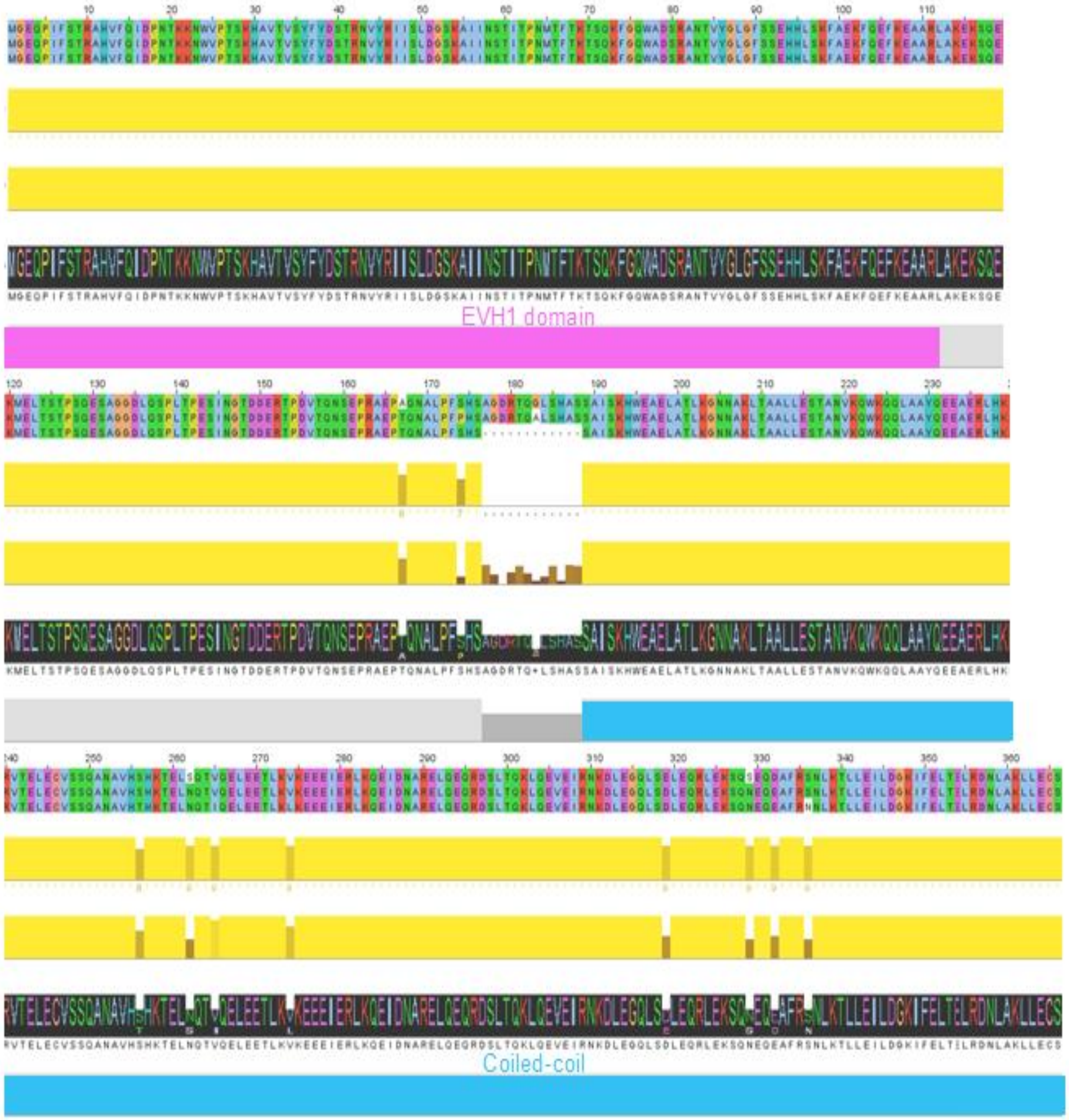


**Figure S2** Multiple sequence alignment of the Human (Q86YM7), Mouse (Q9Z2Y3) and Rat (Q9Z214) Homer1 sequences. Conservation (yellow top), quality (yellow bottom) scores and the consensus sequence (black), along with structural annotation are depicted below the alignment. The canonical isoforms available in UniProt as of February 2025 were used. Alignment and visualization were performed using Jalview.

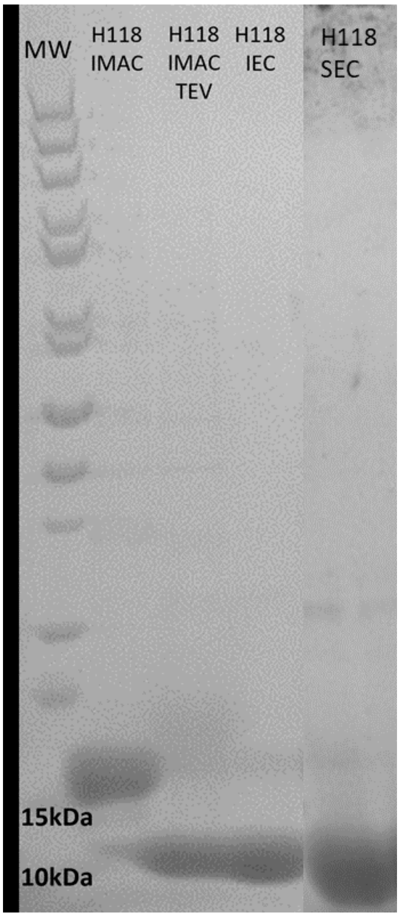


**Figure S3** SDS-PAGE showing the purification steps of the EVH1 domain. The TEV-cleaved form migrates at 13 kDa as expected.

Abbreviations: IMAC - Immobilized Metal Affinity Chromatography, IEC - Ion-Exchange Chromatography, SEC - Size Exclusion Chromatography

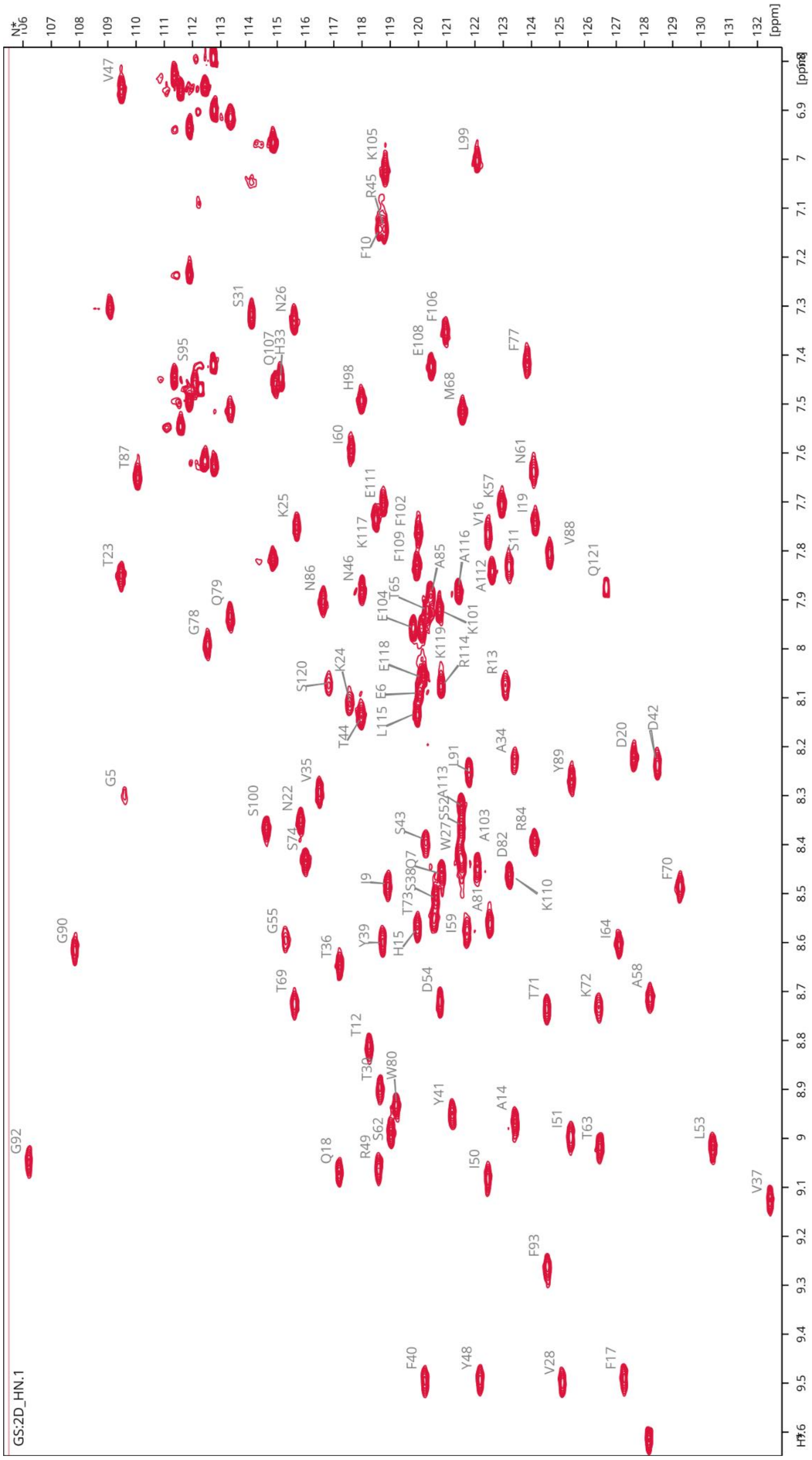


**Figure S4** $^{1}H$-$^{15}N$ HSQC spectrum of EVH1 exhibits signal dispersion characteristic of globular protein. Peak labels show the assigned residues.

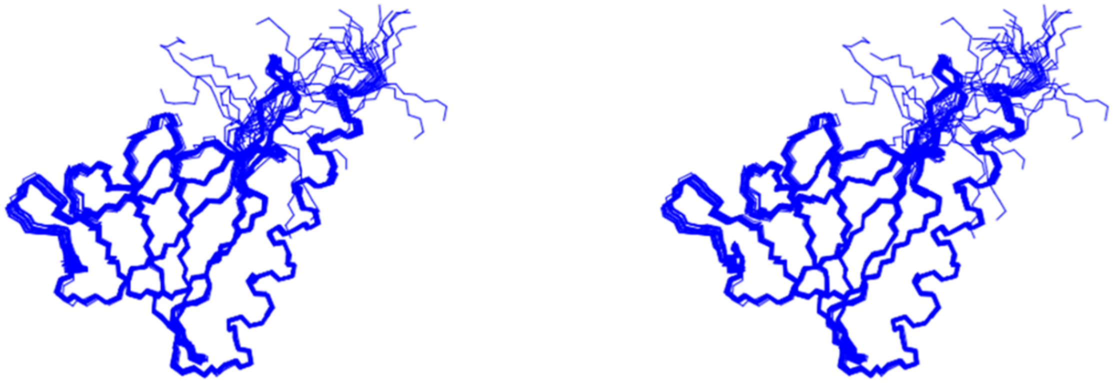

**Figure S5** Stereo view of the structures calculated for the EVH1 domain

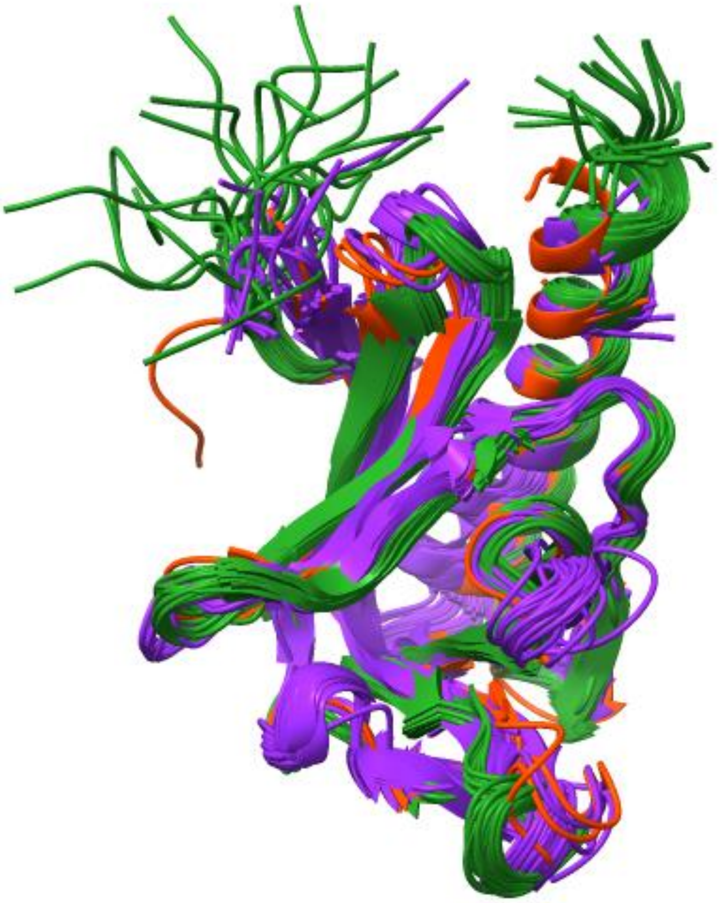

**Figure S6** Structural comparison of the EVH1 domains. Green: the NMR ensemble described in the present study. Orange: other available Homer1 EVH1 structures. Purple: additional EVH1 structures, listed in Table S3. To help visual comparison, additional chains were removed and some structures were truncated at their C-terminus. (extended version of Figure 2C)

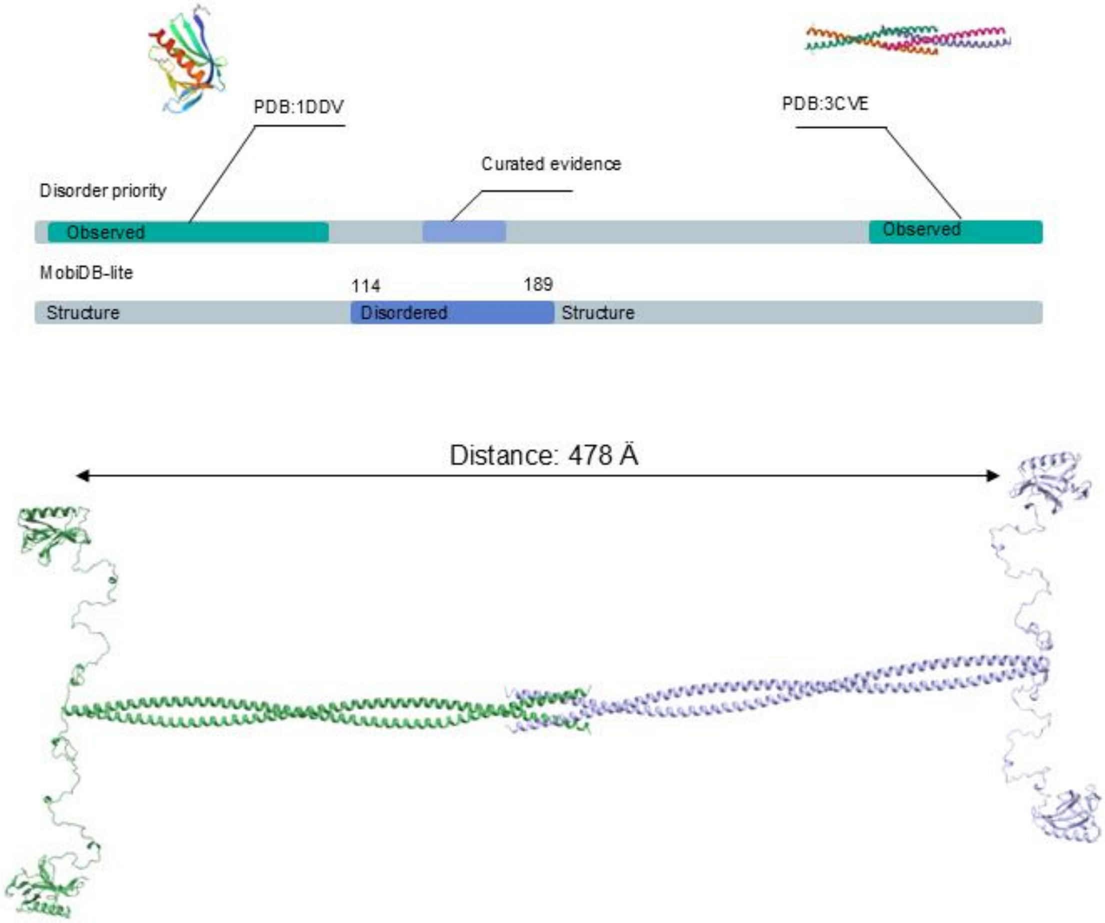


**Figure S7** Modelling of the tetramer coiled-coil. MobiDB was used to determine the boundary between the coiled-coil and the disordered segment. The PDB ids 1ddv and 3cve. denote the experimental structures used for assembling the full model The full length of the coiled-coil in the tetramer spans approximately 478A.

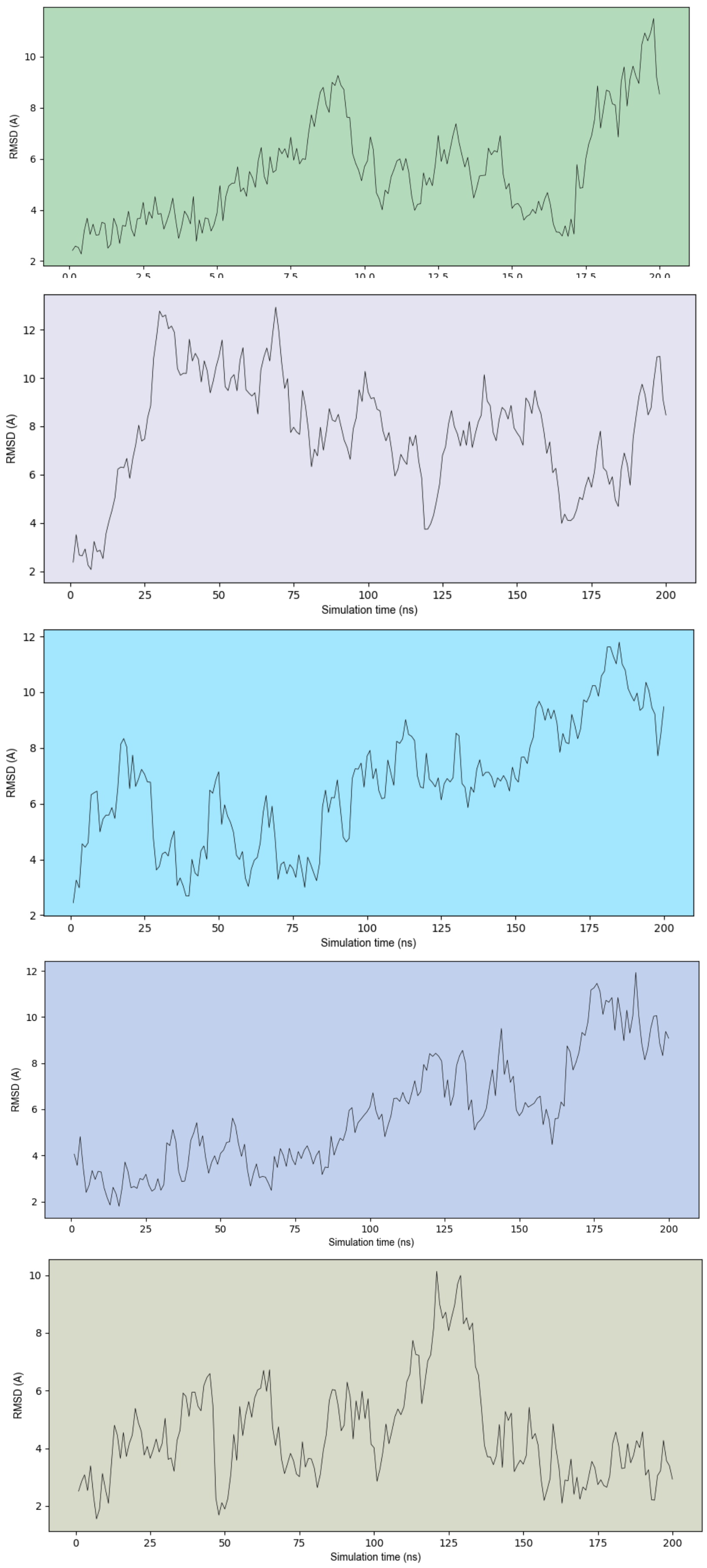


**Figure S8** Global RMSD values calculated for the five parallel 200ns molecular dynamics runs.

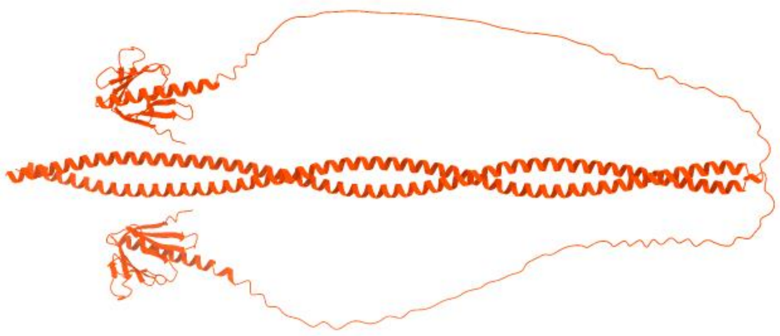

**Figure S9** The predicted model of the dimeric state of Homer1 using AlphaFold2

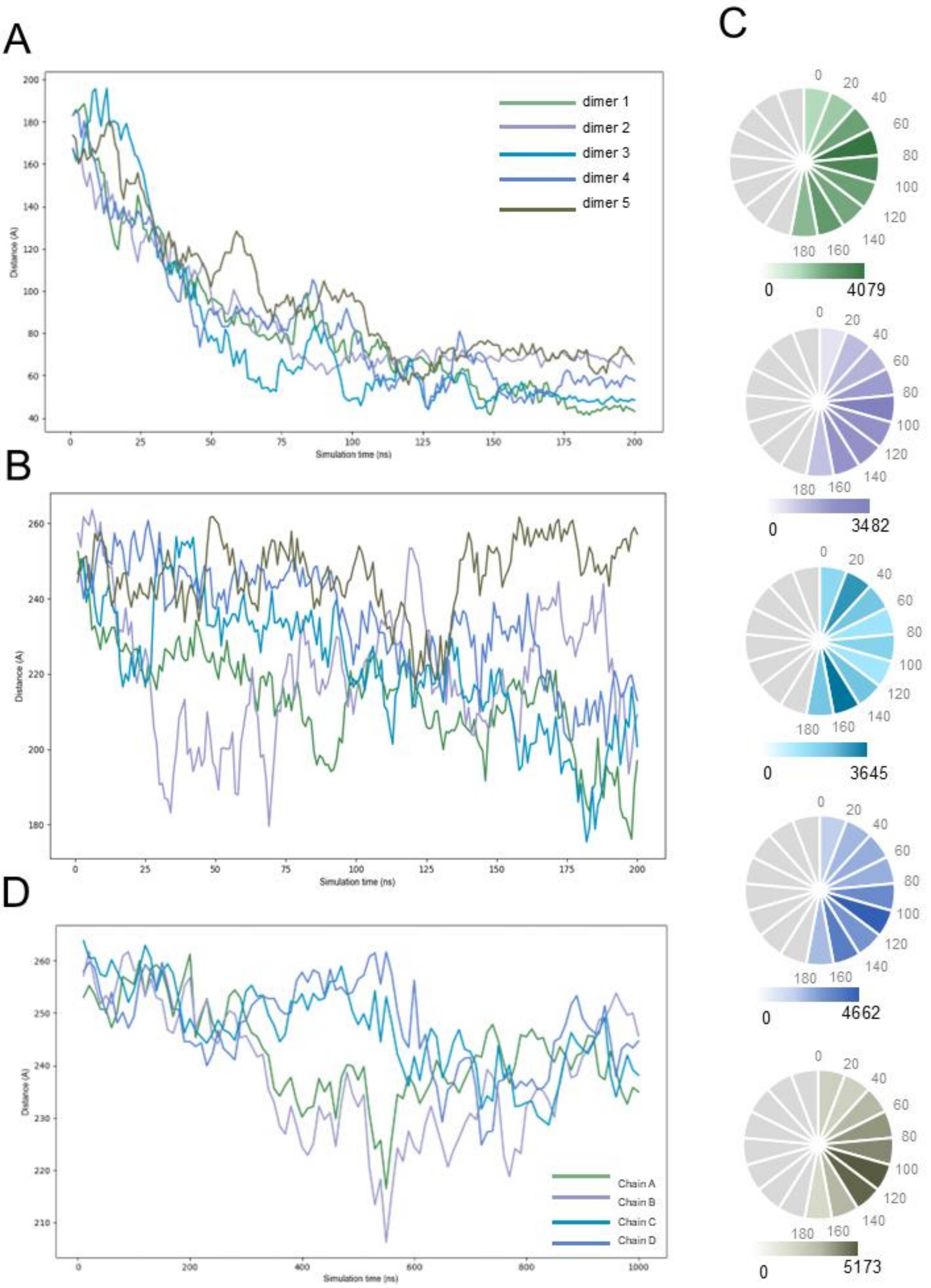
A
B
C
D
dimer 1
dimer 2
dimer 3
dimer 4
dimer 5
Chain A
Chain B
Chain C
Chain D
Distance (A)
Simulation time (ns)
0
4079
0
3482
0
3645
0
4662
0
5173

**Figure S10** Characteristic dynamics of the Homer1 dimer/tetramer.

(A) The average distances of the EVH1 domains between the two chains
(B) Evolution of the kink during the simulation – distance of the extreme N and C terminal CA atoms (Ala 190 and Leu 363) of the coiled-coil
(C) Relative rotation of the two EVH1 domains, calculated by comparing the vectors aligned to beta strand 3.

(D) Evolution of the kink during the simulation – distance of the extreme N and C terminals of the coiled-coil in the tetrameric form

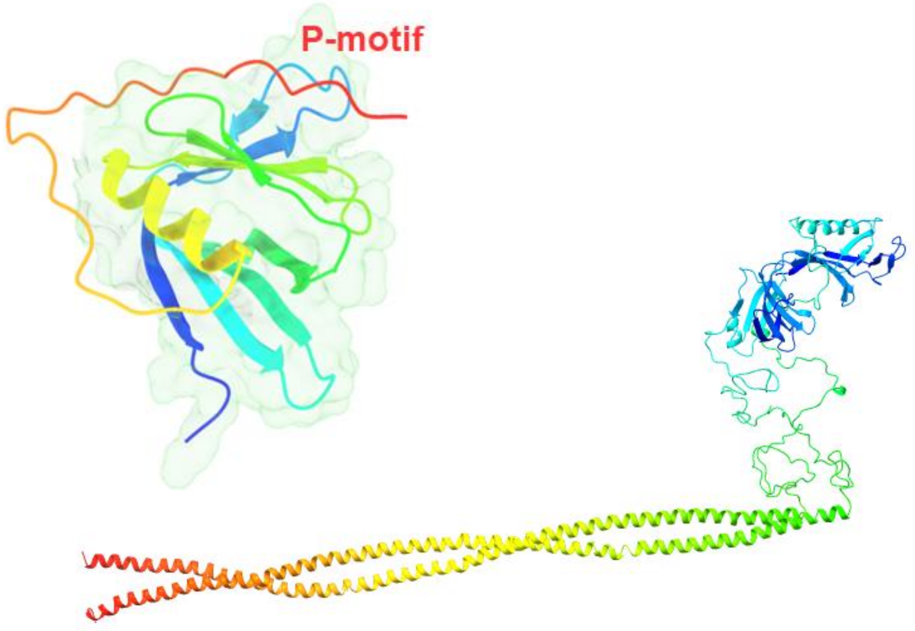


**Figure S11** Intra- and interchain interaction of the P-motif to the EVH1 domain. Intrachain interaction was modelled based on PDB:1ddv, for the interchain interaction our tetramer model was modified. In both cases Modeller from Chimera was used.
